# Molecularly barcoded Zika virus libraries to probe *in vivo* evolutionary dynamics

**DOI:** 10.1101/221564

**Authors:** Matthew T. Aliota, Dawn M. Dudley, Christina M. Newman, James Weger-Lucarelli, Laurel M. Stewart, Michelle R. Koenig, Meghan E. Breitbach, Andrea M. Weiler, Matthew R. Semler, Gabrielle L. Barry, Katie R. Zarbock, Mariel S. Mohns, Emma L. Mohr, Vanessa Venturi, Nancy Schultz-Darken, Eric Peterson, Wendy Newton, Michele L. Schotzko, Heather A. Simmons, Andres Mejia, Jennifer M. Hayes, Saverio Capuano, Miles P. Davenport, Thomas C. Friedrich, Gregory D. Ebel, Shelby L. O’Connor, David H. O’Connor

## Abstract

Defining the complex dynamics of Zika virus (ZIKV) infection in pregnancy and during transmission between vertebrate hosts and mosquito vectors is critical for a thorough understanding of viral transmission, pathogenesis, immune evasion, and potential reservoir establishment. Within-host viral diversity in ZIKV infection is low, which makes it difficult to evaluate infection dynamics. To overcome this biological hurdle, we constructed a molecularly barcoded ZIKV. This virus stock consists of a “synthetic swarm” whose members are genetically identical except for a run of eight consecutive degenerate codons, which creates approximately 64,000 theoretical nucleotide combinations that all encode the same amino acids. Deep sequencing this region of the ZIKV genome enables counting of individual barcode clonotypes to quantify the number and relative proportions of viral lineages present within a host. Here we used these molecularly barcoded ZIKV variants to study the dynamics of ZIKV infection in pregnant and non-pregnant macaques as well as during mosquito infection/transmission. The barcoded virus had no discernible fitness defects *in vivo*, and the proportions of individual barcoded virus templates remained stable throughout the duration of acute plasma viremia. ZIKV RNA also was detected in maternal plasma from a pregnant animal infected with barcoded virus for 64 days. The complexity of the virus population declined precipitously 8 days following infection of the dam, consistent with the timing of typical resolution of ZIKV in non-pregnant macaques, and remained low for the subsequent duration of viremia. Our approach showed that synthetic swarm viruses can be used to probe the composition of ZIKV populations over time *in vivo* to understand vertical transmission, persistent reservoirs, bottlenecks, and evolutionary dynamics.

**Author summary:** Understanding the complex dynamics of Zika virus (ZIKV) infection during pregnancy and during transmission to and from vertebrate host and mosquito vector is critical for a thorough understanding of viral transmission, pathogenesis, immune evasion, and reservoir establishment. We sought to develop a virus model system for use in nonhuman primates and mosquitoes that allows for the genetic discrimination of molecularly cloned viruses. This “synthetic swarm” of viruses incorporates a molecular barcode that allows for tracking and monitoring individual viral lineages during infection. Here we infected rhesus macaques with this virus to study the dynamics of ZIKV infection in nonhuman primates as well as during mosquito infection/transmission. We found that the proportions of individual barcoded viruses remained relatively stable during acute infection in pregnant and nonpregnant animals. However, in a pregnant animal, the complexity of the virus population declined precipitously 8 days following infection, consistent with the timing of typical resolution of ZIKV in non-pregnant macaques, and remained low for the subsequent duration of viremia.

## Introduction

Zika virus (ZIKV; *Flaviviridae, Flavivirus)* infection during pregnancy can cause congenital Zika syndrome (CZS)—a collection of neurological, visual, auditory, and developmental birth defects—in at least 5% of babies [1]. The frequency of vertical transmission is not known, although data suggest that it may be very common, especially if infection occurs during the first trimester [2]. For both pregnant and nonpregnant women, it was previously thought that ZIKV caused an acute self-limiting infection that was resolved in a matter of days. It is now clear that ZIKV can persist for months in other body tissues after it is no longer detectable in blood and in the absence of clinical symptoms [2–7]. During pregnancy, unusually prolonged maternal viremia has been noted, with viral RNA detected in maternal blood up to 107 days after symptom onset [8–11]. The source of virus responsible for prolonged viremia is not known, though it has been speculated that this residual plasma viremia could represent virus release from maternal tissues, the placenta, and/or the fetus.

Recently, we established Indian-origin rhesus macaques *(Macaca mulatta)* as a relevant animal model to understand ZIKV infection during pregnancy, demonstrating that ZIKV can be detected in plasma, CSF, urine, and saliva. In nonpregnant animals viremia was essentially resolved by 10 days post infection [12,13]. In contrast, in pregnant monkeys infected in either the first or third trimester of pregnancy, viremia was prolonged, and was associated with decreased head growth velocity and consistent vertical transmission [2]. Strikingly, significant ocular pathology was noted in fetuses of dams infected with French Polynesian ZIKV during the first trimester [2]. We also showed that viral loads were prolonged in pregnant macaques despite robust maternal antibodies [2]. We therefore aimed to better understand the *in vivo* replication and evolutionary dynamics of ZIKV infection in this relevant animal model.

To do this, we developed a novel “synthetic swarm” virus based on a pathogenic molecular ZIKV clone that allows for tracking and monitoring of individual viral lineages. The synthetic swarm consists of viruses that are genetically identical except for a run of 8 consecutive degenerate nucleotides present in up to ~64,000 theoretical combinations that all encode the same amino acid sequence. This novel barcoded virus is replication competent *in vitro* and *in vivo*, and the number and relative proportion of each clonotype can be characterized by deep sequencing to determine if the population composition changes among or within hosts. Here we demonstrate that this system will provide a useful tool to study the complexity of ZIKV populations within and among hosts; for example, this system can assess bottlenecks following various types of transmission and determine whether non-sterilizing prophylaxis and therapeutics impact the composition of the virus population. Moreover, data from molecularly barcoded viruses will help inform research of ZIKV infection during pregnancy by providing a better understanding of the kinetics of tissue reservoir establishment, maintenance, and reseeding.

## Results

### Generation and characterization of a molecularly barcoded virus stock

Molecular barcoding has been a useful tool to study viruses including simian immunodeficiency virus, influenza virus, poliovirus, Venezuelan equine encephalitis virus, and West Nile virus, establishing conceptual precedent for our approach [14–20]. To generate barcoded ZIKV, we introduced a run of eight consecutive degenerate codons into a region of NS2A (amino acids 144–151) that allows for every possible synonymous mutation to occur in the ZIKV infectious molecular clone (ZIKV-IC) derived from the Puerto Rican isolate ZIKV-PRVABC59 [21]. Following bacteria-free cloning and rolling circle amplification (RCA), linearized and purified RCA reaction products were used for virus production via transfection of Vero cells. All produced virus was collected, pooled, and aliquoted into single-use aliquots, such that single aliquots contain a representative sampling of all genetic variants generated; this barcoded synthetic swarm virus was termed ZIKV-BC-1.0.

We used a multiplex-PCR approach to deep sequence the entire coding genome of the ZIKV-BC-1.0 stock, as well as the ZIKV-IC from which ZIKV-BC-1.0 was derived. For each stock, 1 x 10^6^ viral RNA templates were used in each cDNA synthesis reaction (**Table 1**), and both stocks were sequenced in duplicate. We identified two nucleotide positions outside of the barcode region that encoded fixed differences between ZIKV-IC and ZIKV-BC-1.0, when compared to the KU501215 reference that we used for mapping. The variant at site 1964 encodes a nonsynonymous change (V to L) in Envelope, and the variant at site 8488 encodes a synonymous substitution in NS5. The variant at site 1964 was also present in our ZIKV-PRVABC59 stock (see [22]), and Genbank contains records for two sequences that match this sequence (accession numbers KX087101 and KX601168) and two that do not (KU501215 and KX377337). In addition, a single nucleotide position in NS5 contained an 80/20 ratio of C-to-T nucleotide substitutions in ZIKV-BC-1.0, but was fixed as a C in ZIKV-IC. The C-to-T change is a synonymous mutation in a leucine codon. There were no other high-frequency variants that differentiate the two stocks outside of the barcode region in the remainder of the genome encoding the polyprotein open reading frame.

**Table 1.** Number of viral templates used to characterize the full genome sequences of the two ZIKV stocks.

| Sample | Number of vRNA templates put into the cDNA synthesis reaction | Number of cDNA templates put into the PCR reaction* |
| --- | --- | --- |
| ZIKV-IC stock | $1 \times 10^6$ | 119,048 |
| ZIKV-BC-1.0 | $1 \times 10^6$ | 119,048 |

The number of cDNA templates was calculated based on the number of vRNA templates that were put into the cDNA synthesis reaction, and then the amount of cDNA that was used for the PCR reaction.

### Diversity of barcode sequences in the stock of ZIKV-BC-1.0

We then characterized the diversity of barcode sequences present in the ZIKV-BC-1.0 stock prior to *in vitro* and *in vivo* studies. To exclude PCR artifacts that could have biased our barcode estimates, we identified every distinct barcode sequence in either the barcoded stock (ZIKV-BC-1.0) or in the non-barcoded parental infectious clone (ZIKV-IC) in the region of NS2A encompassing the barcode. We then calculated the frequency of each of these barcode sequences in the replicates of both stocks. Many of the sequences were detected in only one or two of the replicates, so the frequency of some of the sequences was actually zero in one replicate, even if it was detected in a second replicate. We then calculated the arithmetic mean (0.0018%) and standard deviation (0.015%) of the frequencies of all the non-wild type sequences in the 24 nucleotide region in ZIKV-IC that corresponds to the barcoded 24 nucleotides in ZIKV-BC-1.0. We used the mean and standard deviation frequencies from ZIKV-IC to calculate the “noisiness” inherent in deep sequencing the barcoded region. Authentic barcodes were defined as those whose frequency was greater than three standard deviations higher than this threshold; by this standard the minimum frequency for an authentic barcode sequence in ZIKV-BC-1.0 was 0.047% in at least one of the two replicates. Using this criterion, we detected 57 distinct barcodes in ZIKV-BC-1.0. Of the 57 barcodes, 20 were detected in both sequencing replicates of the ZIKV-BC-1.0 stock at a frequency of 0.5% or greater (present in >168 sequencing reads) and were given independent labels (e.g. Zika_BC01, Zika_BC02, etc.) to simplify reporting. Barcodes 21 to 57 were also tracked during infection, but then categorized as ‘Other barcodes’ in the graphs shown throughout. Any other barcodes detected below the 0.05% threshold were categorized as ‘Noise’.

To ascertain whether input RNA template numbers influence barcode composition, we sequenced a dilution series of viral RNA templates in triplicate (**Fig 1, Table 2, and Table S1**). When we used 10,000 input vRNA templates, we enumerated an average of 55 of the 57 barcodes. At 2000 and 500 input vRNA templates, we enumerated an average of 46 and 33 of the 57 barcodes, respectively. For 250, 100, and 50 input templates, the average number of enumerated barcodes was 26.7 ± 3.9, indicating that the number of unique barcodes we enumerated was consistent between 50 and 250 input vRNA templates. It is also important to note that an average of 1.46% of the sequences in the 50 input vRNA template samples were considered ‘noise’ because they contained barcodes that were not among the 57 we enumerated from the ZIKV-BC-1.0 stock, while an average of 3.56% of the sequences in the 10,000 input vRNA template samples were considered ‘noise.’ This observation suggests that, on average, while we sequence fewer unique authentic templates when fewer input molecules are used, the reduction of input molecules does not increase the detection of spurious, ‘noise’ barcodes.

**Figure 1.**
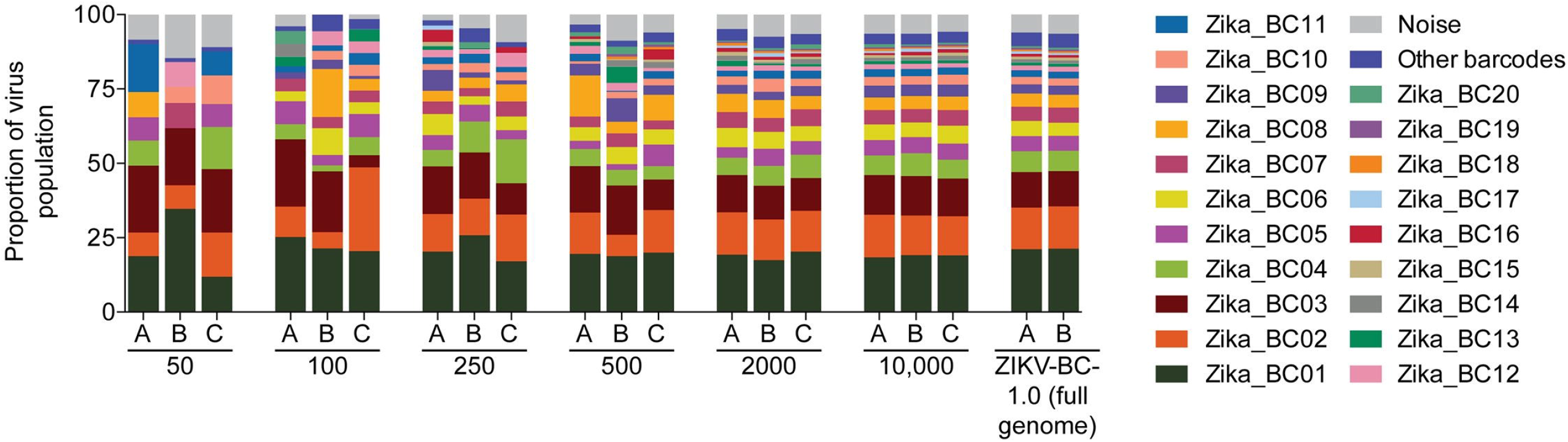
Sequencing of the titrated ZIKV-BC-1.0 stock. Dilutions of ZIKV-BC-1.0 were reverse-transcribed, PCR-amplified with a single primer pair, and sequenced, as described in the materials and methods. The number of vRNA templates that were used for each cDNA synthesis reaction is shown below the graph. The theoretical number of cDNA molecules used in each PCR reaction is shown in Table 2. Each dilution was sequenced in triplicate, with individual replicates labeled A, B, and C, the frequency of each barcode was enumerated and are shown. ‘Other barcodes’ were the barcodes present in the list of the Top 57. ‘Noise’ represents sequences detected in the barcode region that did not match the Top 57 barcode sequences.

**Table 2.** Number of viral templates used in the titration analysis of ZIKV-BC-1.0

| Sample | Number of vRNA templates put into the cDNA synthesis reaction | Number of theoretical cDNA templates put into the PCR reaction* |
| --- | --- | --- |
| 50 copies | 50 | 30 |
| 100 copies | 100 | 60 |
| 250 copies | 250 | 149 |
| 500 copies | 500 | 298 |
| 2000 copies | 2000 | 1,190 |
| 10,000 copies | 10,000 | 5,952 |

The number of cDNA templates was calculated based on the number of vRNA templates that were put into the cDNA synthesis reaction, and then the amount of cDNA that was used for the PCR reaction.

We also examined diversity and similarity across sequencing replicates in this titration experiment using all the detected sequences, including the ‘noise.’ Not surprisingly, Simpson’s diversity increased when a greater number of input templates were used, plateauing at 500 input copies (**Fig S1**). When comparing similarity across replicates, the samples with 2,000 and 10,000 inputs had the highest Morisita-Horn similarity index (**Fig S2**). Unfortunately, it is not possible to obtain a large number of input templates at all timepoints from ZIKV-infected pregnant animals. The detection of barcodes at high frequency and ‘noise’ at low frequency when using low template input suggests that the detection of a barcode in these samples is likely believable (**Fig S3**). It is important to note, however, the absence of a barcode in sequencing reads from a particular experiment could mean that either the barcode was not present at that timepoint or that it was present in the biological sample but not at a high enough concentration to be detected when sequencing from a small number of templates.

### Molecularly-barcoded ZIKV *in vivo* replication kinetics and barcode dynamics

Prior to use in nonhuman primates, viral infectivity and replication of ZIKV-BC-1.0 was assessed *in vitro* using Vero, LLC-MK2, C6/36, and Aag2 cells. Viral growth curves were similar between ZIKV-BC-1.0, infectious clone-derived virus (ZIKV-IC), and wild-type ZIKV-PRVABC59 (ZIKV-PR) (**See Weger-Lucarelli et al. concomitant submission**). These results demonstrated that the insertion of degenerate nucleotides in the barcode viral genome did not have a measurable deleterious effect on either infectivity or replicative capacity *in vitro*. To confirm that ZIKV-BC-1.0 did not have any replication defects *in vivo*, we assessed its replication capacity in rhesus macaques. Three rhesus macaques were inoculated subcutaneously with 1 x 10^4^ PFU of ZIKV-BC-1.0. All three animals were productively infected with ZIKV-BC-1.0, with detectable plasma viral loads one day post inoculation (dpi) (**Fig 2**). Plasma viral loads in all three animals peaked between two and four dpi, and ranged from 2.34 x 10^3^ to 9.77 x 10^4^ vRNA copies/mL. Indeed, ZIKV-BC-1.0 displayed viral replication kinetics comparable to ZIKV-IC and ZIKV-PR (i.e., area under the curve was not significantly different (Student’s *t*-test), p=0.355 and 0.229, respectively); and replication kinetics were comparable to previous studies with other strains of ZIKV in nonpregnant rhesus macaques [12,13,23].

**Figure 2.**
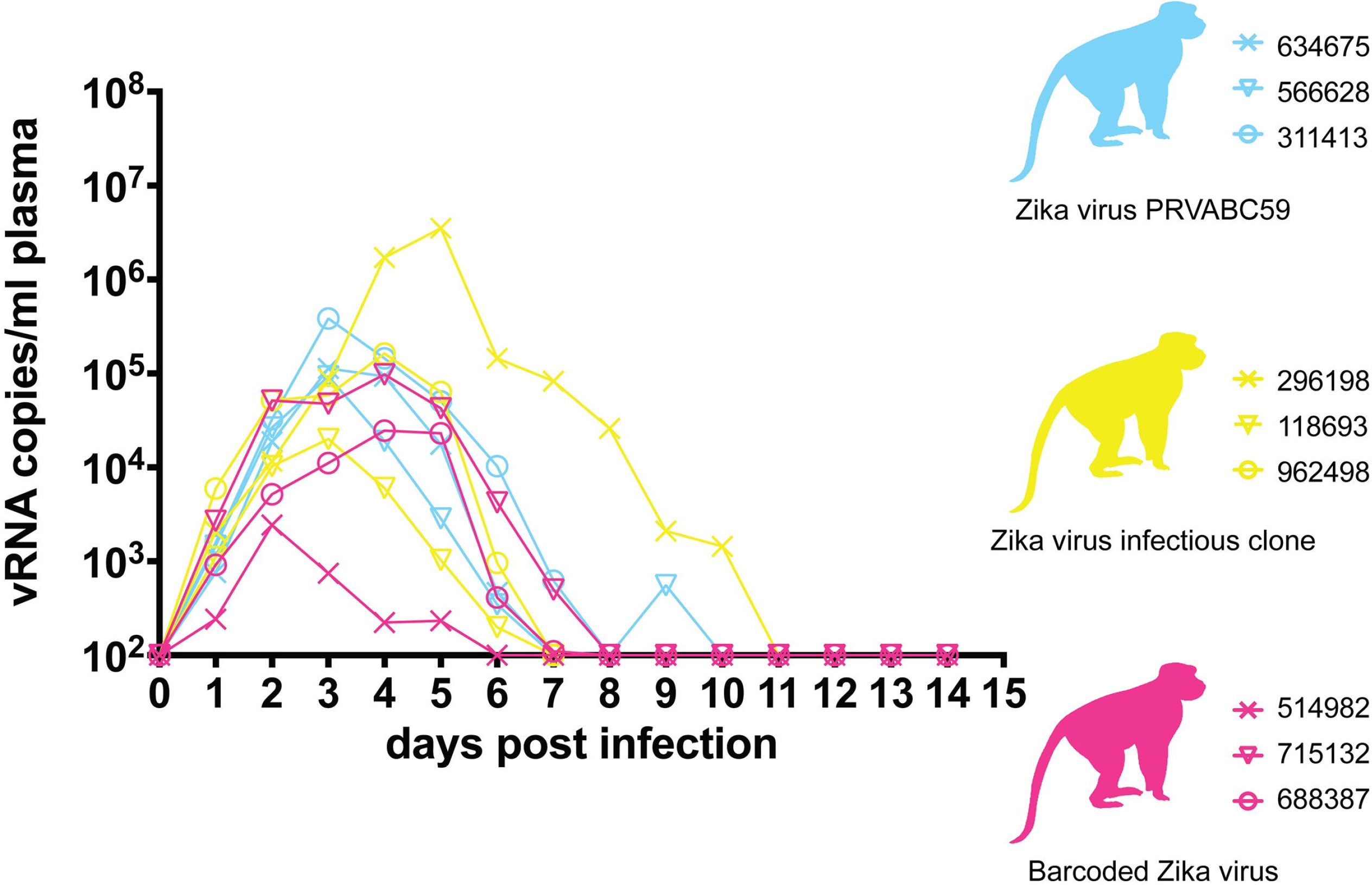
Longitudinal detection of Zika vRNA in plasma from animals inoculated with ZIKV-PR (blue), ZIKV-IC (yellow), or ZIKV-BC-1.0 (magenta). Zika vRNA copies per ml blood plasma. The y-axis crosses the x-axis at the limit of quantification of the qRT-PCR assay (100 vRNA copies/ml).

We also infected a single pregnant macaque (776301) by subcutaneous inoculation of 1 x 10^4^ PFU of ZIKV-BC-1.0. This animal had been exposed to dengue virus serotype 3 (DENV-3; strain Sleman/78) approximately one year prior to inoculation with ZIKV-BC-1.0. To evaluate cross-reactive neutralizing antibody (nAb) responses elicited by prior exposure to DENV-3 in this animal, serum was obtained prior to inoculation with ZIKV-BC-1.0. Neutralization curves with both DENV-3 and ZIKV revealed that DENV-3 immune sera did not cross-react with ZIKV, whereas DENV-3 was potently neutralized (**Fig 3A**). The animal then was infected with ZIKV-BC-1.0 at 35 days of gestation (mid-first trimester; rhesus term is 165 ± 10 days) and had detectable plasma viral loads for 64 dpi (Fig 3B); consistent with replication kinetics of wildtype ZIKV in both pregnant macaques [2] and humans [8,9,24]. The animal also had four days of detectable vRNA in urine but no detectable vRNA (**Fig 3B**) in the amniotic fluid on 22, 36, 50, or 120 dpi (57, 71, 85, 155 days gestation, respectively). By 29 dpi neutralization curves of both viruses revealed a similar profile, indicating the production of a robust maternal nAb response to ZIKV (**Fig 3A**) coincident with prolonged plasma viral loads, similar to what has been shown previously in other ZIKV-infected pregnant macaques [2]. DENV-3 neutralization curves at 0 and 29 dpi were indistinguishable (Fig 3A).

**Figure 3.**
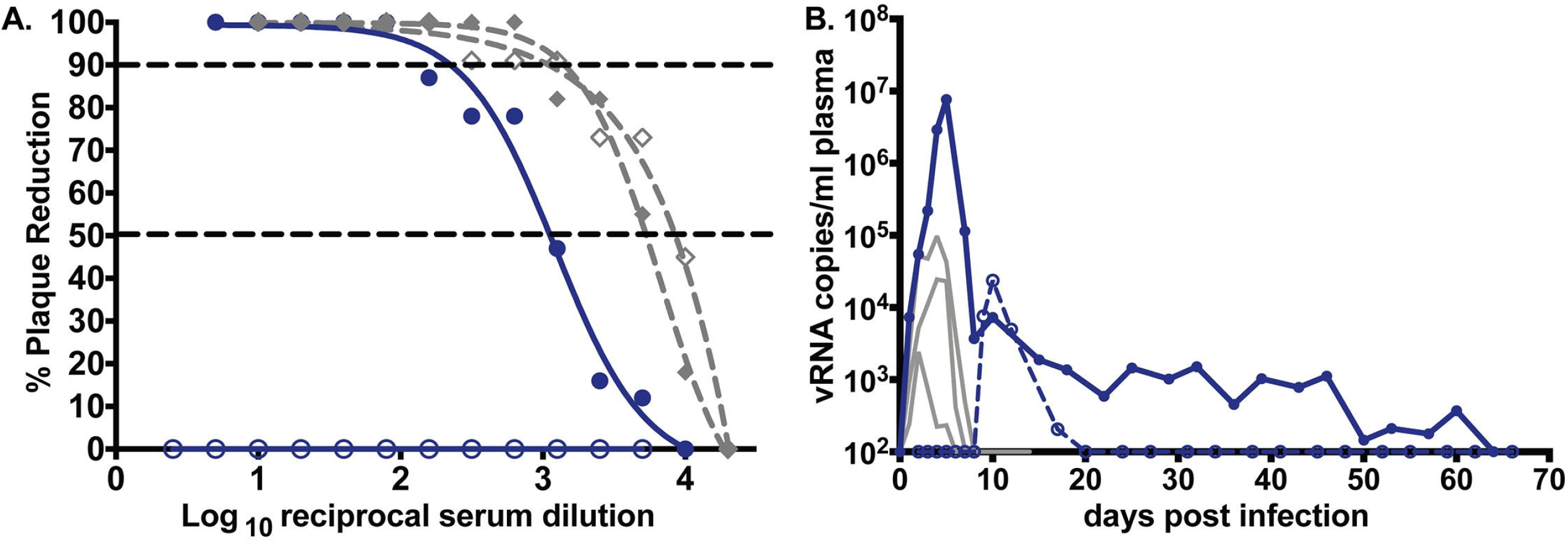
Maternal Zika vRNA loads and neutralization curves. A.) Neutralization by ZIKV and DENV-3 immune sera from a pregnant ZIKV-BC-1.0-infected macaque. Immune sera from a macaque infected with ZIKV-BC-1.0 during the first trimester of pregnancy was tested for its capacity to neutralize DENV-3 (gray dashes) and ZIKV-PR (blue). Infection was measured by plaque reduction neutralization test (PRNT) and is expressed relative to the infectivity of ZIKV-PR in the absence of serum. The concentration of sera indicated on the x-axis is expressed as log10 (dilution factor of serum). The EC90 and EC50, estimated by non-linear regression analysis, are also indicated by a dashed line. Neutralization curves for each virus (ZIKV, solid blue; DENV-3, dashed grey) at 0 (open symbols) and 28 (closed symbols) dpi are shown. **B.**) Zika vRNA copies per ml blood plasma (solid lines) or urine (dashed line). Blue tracings represent the animal infected with ZIKV-BC-1.0 at 35 days gestation. The day of gestation is estimated +/− 2 days. Grey tracings represent viremia in nonpregnant/male rhesus monkeys infected with the identical dose of ZIKV-BC-1.0 (Figure 2). The y-axis crosses the x-axis at the limit of quantification of the qRT-PCR assay (100 vRNA copies/ml).

The pregnancy progressed without adverse outcomes, and at 155 days of gestation, the fetus was surgically delivered, euthanized, and tissues collected. The fetus had no evidence of microcephaly or other abnormalities upon gross examination. Approximately 60 fetal and maternal tissues (see **Table 3** for a complete list) were collected sterilely for histopathology and vRNA by QRT-PCR. No ZIKV RNA was detected in any samples collected from the fetus, suggesting that vertical transmission did not occur. This was surprising, as from seven neonatal macaques we have examined to date, this was the only animal found not to have detectable ZIKV RNA in tissues. This also shows that prolonged maternal viremia can be uncoupled from detection of ZIKV RNA in fetal tissues at birth. Fetal histology revealed normal CNS anatomy, minimal to mild suppurative lymphadenitis of the inguinal lymph node, minimal multifocal lymphocytic deciduitis, and mild multifocal placental infarction with suppurative villositis, similar to changes noted in previous *in utero* ZIKV infections [2]. These data demonstrate that ZIKV-BC-1.0 is fully functional *in vivo* with replication kinetics indistinguishable from other ZIKV strains and suggest that the inclusion of the barcode did not impair infectivity or replication in adult macaques.

**Table 3.** Complete list of tissues examined for ZIKV-BC-1.0 RNA from 776301 and her fetus.

|  | **Dam** | **Fetus** |
| --- | --- | --- |
| mesenteric LN | - | ND |
| spleen | - | ND |
| adipose tissue | ND | - |
| adrenal gland | ND | - |
| amniotic/chorionic membrane | ND | - |
| aorta-thoracic | ND | - |
| articular-cartilage | ND | - |
| axillary LN | ND | - |
| bile aspirate | ND | - |
| bone marrow | ND | - |
| cerebrum (9 sections) | ND | - |
| cervical spinal cord | ND | - |
| colon | ND | - |
| cord blood-serum | ND | - |
| cornea | ND | - |
| decidua | ND | - |
| dura mater | ND | - |
| epidermis/dermis abdomen | ND | - |
| esophagus | ND | - |
| eye-aqueous humor | ND | - |
| femur bone | ND | - |
| heart | ND | - |
| inguinal LN | ND | - |
| jejunum | ND | - |
| kidney | ND | - |
| liver | ND | - |
| lumbar spinal cord | ND | - |
| lung | ND | - |
| meconium | ND | - |
| mesenteric LN | ND | - |
| muscle-quadriceps | ND | - |
| optic nerve | ND | - |
| ovary | ND | - |
| pancreas | ND | - |
| pericardium | ND | - |
| pituitary gland | ND | - |
| placental disc 1 | ND | - |
| placental disc 2 | ND | - |
| retina | ND | - |
| sclera | ND | - |
| spleen | ND | - |
| stomach | ND | - |
| submandibular LN | ND | - |
| terminal blood draw-plasma | ND | - |
| terminal CSF | ND | - |
| thoracic spinal cord | ND | - |
| thymus | ND | - |
| thyroid | ND | - |
| tongue | ND | - |
| tonsil | ND | - |
| tracheobronchial LN | ND | - |
| umbilical cord | ND | - |
| urinary bladder | ND | - |
| urine-aspirate | ND | - |
| uterus | ND | - |
| uterus-placental bed | ND | - |

-, ZIKV RNA below the limit of detection
LN, lymph node
ND, no data.

### Evaluation of barcodes during acute infection of nonpregnant macaques

We deep sequenced the viruses replicating in the nonpregnant animals who were infected with ZIKV-BC-1.0 and ZIKV-IC (**Fig 4A and B, Table 4, and Table S2**). In each group of three animals, we sequenced viruses at two time points from two animals, and then one time point from a third animal. In animals infected with ZIKV-IC, we found that >95% of sequences in the virus stock and all three animals were wild type across the 24 nucleotides that corresponded to where the barcode was located in ZIKV-BC-1.0.

**Figure 4.**
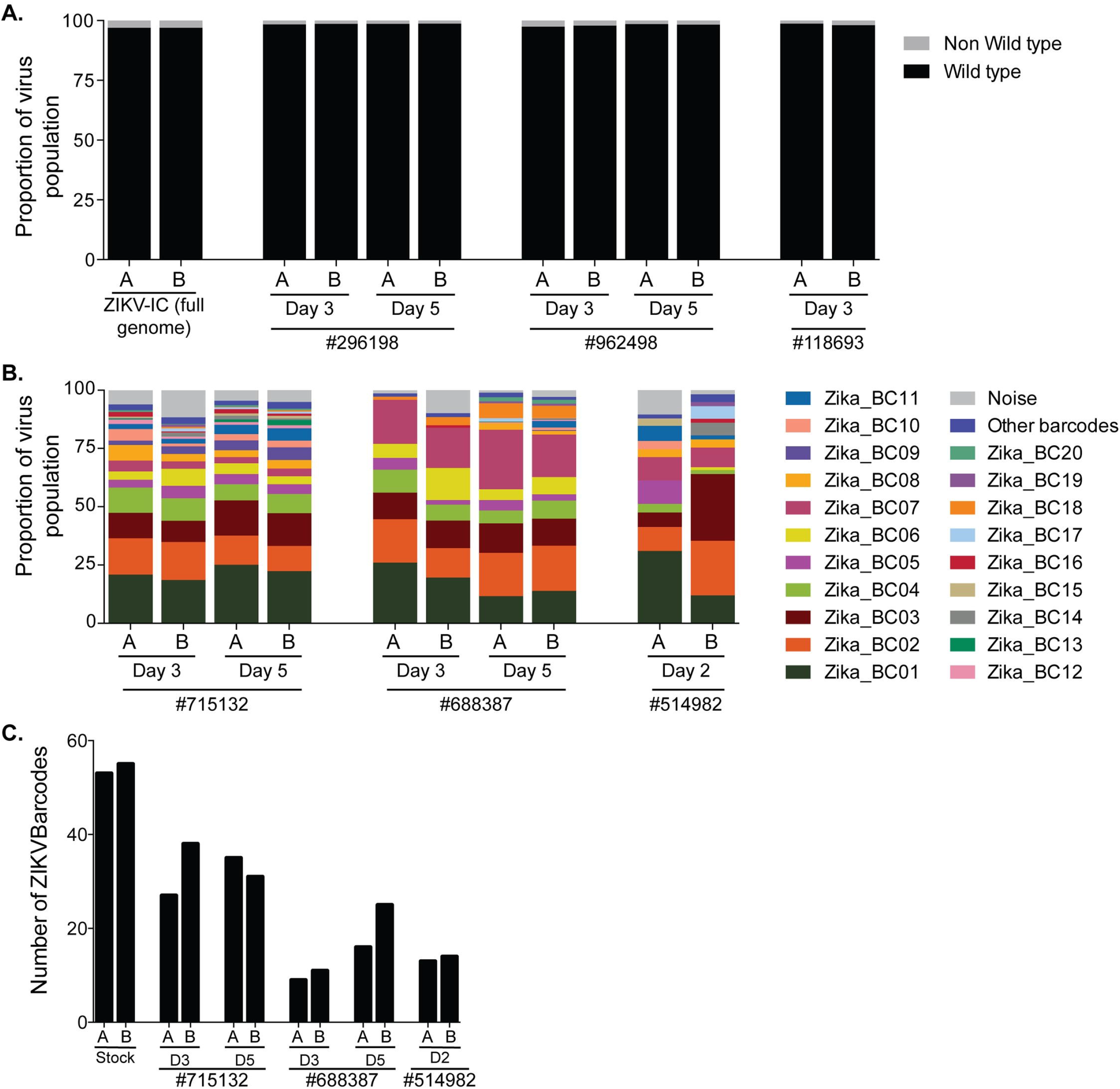
Sequencing of ZIKV-BC-1.0 and ZIKV-IC isolated from nonpregnant rhesus macaques. ZIKV RNA was isolated from plasma at the indicated time points from each of the animals infected with **A.**) ZIKV-IC or **B.**) ZIKV-BC-1.0. Viral RNA was reverse transcribed and then multiplex PCR was performed as described in materials and methods. PCR products were tagged and sequenced. **A.**) The sequence mapping to the region containing the molecular barcode was interrogated for ZIKV-IC, and the frequency of wild type and non-wild type ZIKV sequences are shown. The theoretical number of cDNA molecules used in each PCR reaction is shown in Table 4. Each sample was sequenced in duplicate, as labeled by A and B. **B.**) The frequency of each barcode in the population is shown for ZIKV-BC-1.0. ‘Other barcodes’ were the barcodes present in the list of the Top 57. ‘Noise’ represents sequences detected in the barcode region, but did not match the Top 57. **C.**) The number of barcodes detected at a frequency of greater than 0.047% in the three nonpregnant animals and the stock were counted. The data for each individual replicate are shown. Some barcodes were detected at a frequency of 0.047% or greater in replicate A, but not replicate B, and vice versa.

**Table 4.**
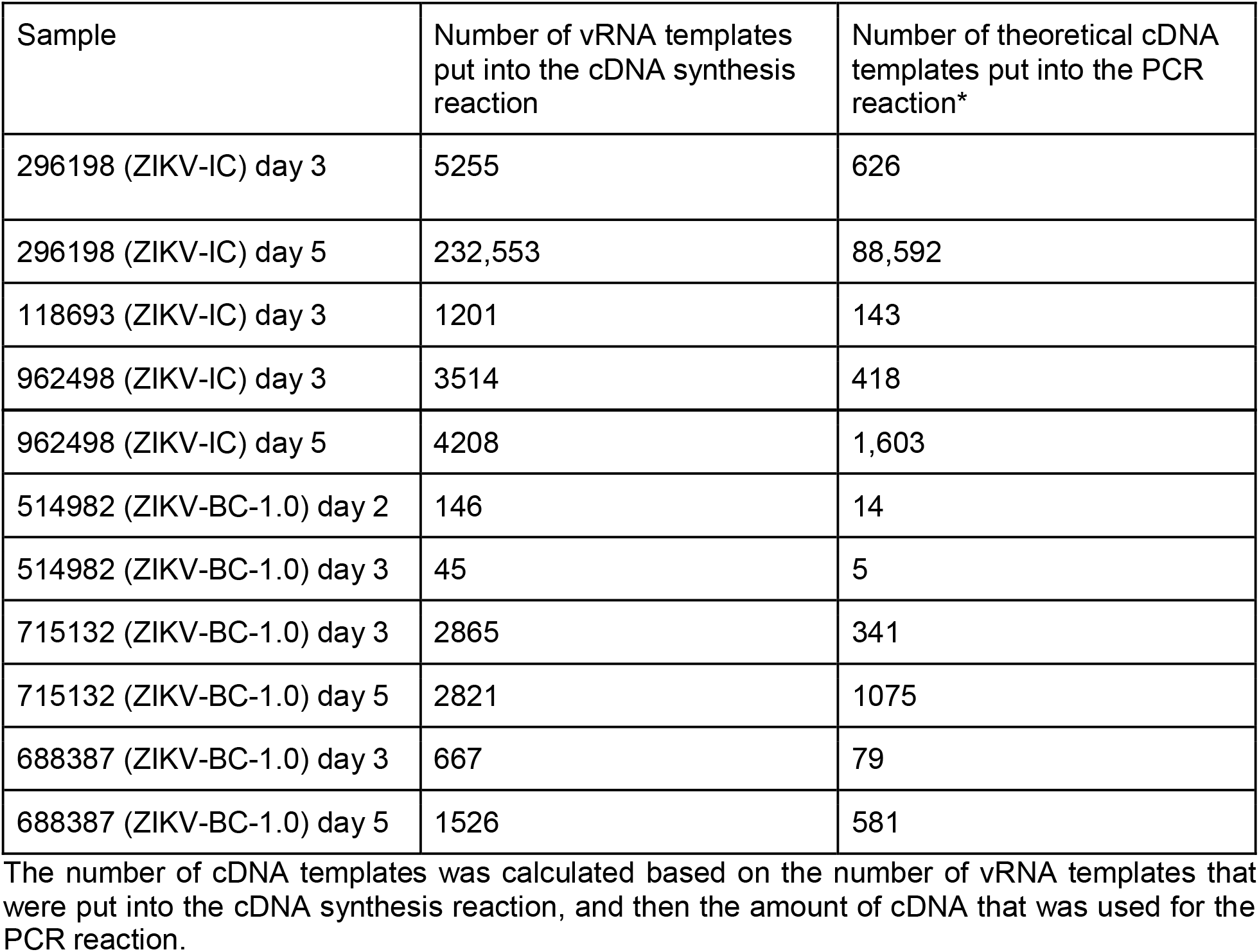
Number of viral templates sequenced from nonpregnant animals infected with ZIKV-IC or ZIKV-BC-1.0

| Sample | Number of vRNA templates put into the cDNA synthesis reaction | Number of theoretical cDNA templates put into the PCR reaction* |
| --- | --- | --- |
| 296198 (ZIKV-IC) day 3 | 5255 | 626 |
| 296198 (ZIKV-IC) day 5 | 232,553 | 88,592 |
| 118693 (ZIKV-IC) day 3 | 1201 | 143 |
| 962498 (ZIKV-IC) day 3 | 3514 | 418 |
| 962498 (ZIKV-IC) day 5 | 4208 | 1,603 |
| 514982 (ZIKV-BC-1.0) day 2 | 146 | 14 |
| 514982 (ZIKV-BC-1.0) day 3 | 45 | 5 |
| 715132 (ZIKV-BC-1.0) day 3 | 2865 | 341 |
| 715132 (ZIKV-BC-1.0) day 5 | 2821 | 1075 |
| 688387 (ZIKV-BC-1.0) day 3 | 667 | 79 |
| 688387 (ZIKV-BC-1.0) day 5 | 1526 | 581 |

The number of cDNA templates was calculated based on the number of vRNA templates that were put into the cDNA synthesis reaction, and then the amount of cDNA that was used for the PCR reaction.

We counted the number of authentic barcodes detected at a frequency of 0.047% or greater in the stock and the plasma of the nonpregnant animals infected with ZIKV-BC-1.0. Whereas we counted a total of 57 barcodes at this threshold in the stock, we found a range of 9 to 38 barcodes in each of the samples from the animals (Fig 4C). We then compared the frequency of the individual barcodes in the plasma of these three animals relative to that in the stock. In animal 715132, the frequency of each barcode at day 3 was maximally 3.4 percentage points different from the frequency of each barcode in the stock. By day 5, the barcode frequencies changed by, at most, 4.6 percentage points when compared to day 3. In animal 688387, we found that the frequency of Zika_BC07 at day 3 in the population was an average of 18.1%, which was markedly greater than the 5% frequency of Zika_BC07 observed in the stock. The frequency of Zika_BC07 continued to increase to 21.9% of the population at day 5. As a result, the frequency of each barcode at day 3 was maximally 13.1 percentage points different from the the frequency of each barcode in the stock, and the barcode frequencies at day 5 were up to to 10.1 percentage points different from day 3. Unfortunately, we were only able to acquire sequence data from animal 514982 at day 2 post infection, but the frequency of each barcode was maximally 5.6 percentage points different from the frequency of each barcode in the stock.

We also examined the sequences outside the barcode region to determine if there were additional nucleotide differences present in the virus population as it replicated in animals. There were small fluctuations in some viral SNPs, but we detected no dramatic shifts in nucleotide frequencies among viruses replicating in vivo, except at site 9581, which is synonymous. In the ZIKV-BC-1.0 stock, there was a mixture of T and C nucleotides (22% and 78% of sequences, respectively) at this site. This position remained a mixture in the animals, but the ratios fluctuated. It dipped to a ratio of 10/90 in animal 688387 at day 5 to as high as 30/70 in animal 715132 at day 5. Overall, there were no new mutations that were detected at greater than 10% frequency in both replicates in the virus populations during the first 5 days after infection in nonpregnant animals.

### Evaluation of barcodes during pregnancy

We also deep sequenced the barcode in virus populations replicating in the one pregnant animal (776301) infected with ZIKV-BC-1.0. Recognizing that the later time points from this animal had persistent, but low plasma viral loads, we modified our sequencing approach to prepare one tube of cDNA, and then split it into two independent PCR reactions that amplified small fragments (131bp and 178bp) spanning the region containing the barcode (**Fig 5A, Table 5, and Table S3**). We quantified the number of authentic barcodes we detected at a frequency of 0.047% or greater (**Fig 5B**). At days 3, 5, and 7, we detected an average of 39.3 ± 2.6 barcodes. This declined precipitously to an average of 9 barcodes at days 8 and 10. After day 10, we did not detect more than 7 barcodes, and, in fact, we only detected 1 authentic barcode present at a frequency of 0.047% or greater at some timepoints. Likewise, barcode diversity, as measured by Simpson’s diversity index, also declined beginning at day 8 and remained low throughout the duration of infection (**Fig 5C**). Interestingly, some barcodes, such as Zika_BC02, were not detected at later timepoints, even though it had been present at ~15% during early infection. Other barcodes, such as Zika_BC07, 08, and 09, became more common at later time points, even though they were only present at ~2-5% during early infection. Unfortunately, with such low virus input templates at the late time points, there were differences between replicates indicative of sampling uncertainty. With the exception of two samples (day57_A and day60_B), however, greater than 95% of the sequences matched one of the 57 authentic barcodes.

**Figure 5.**
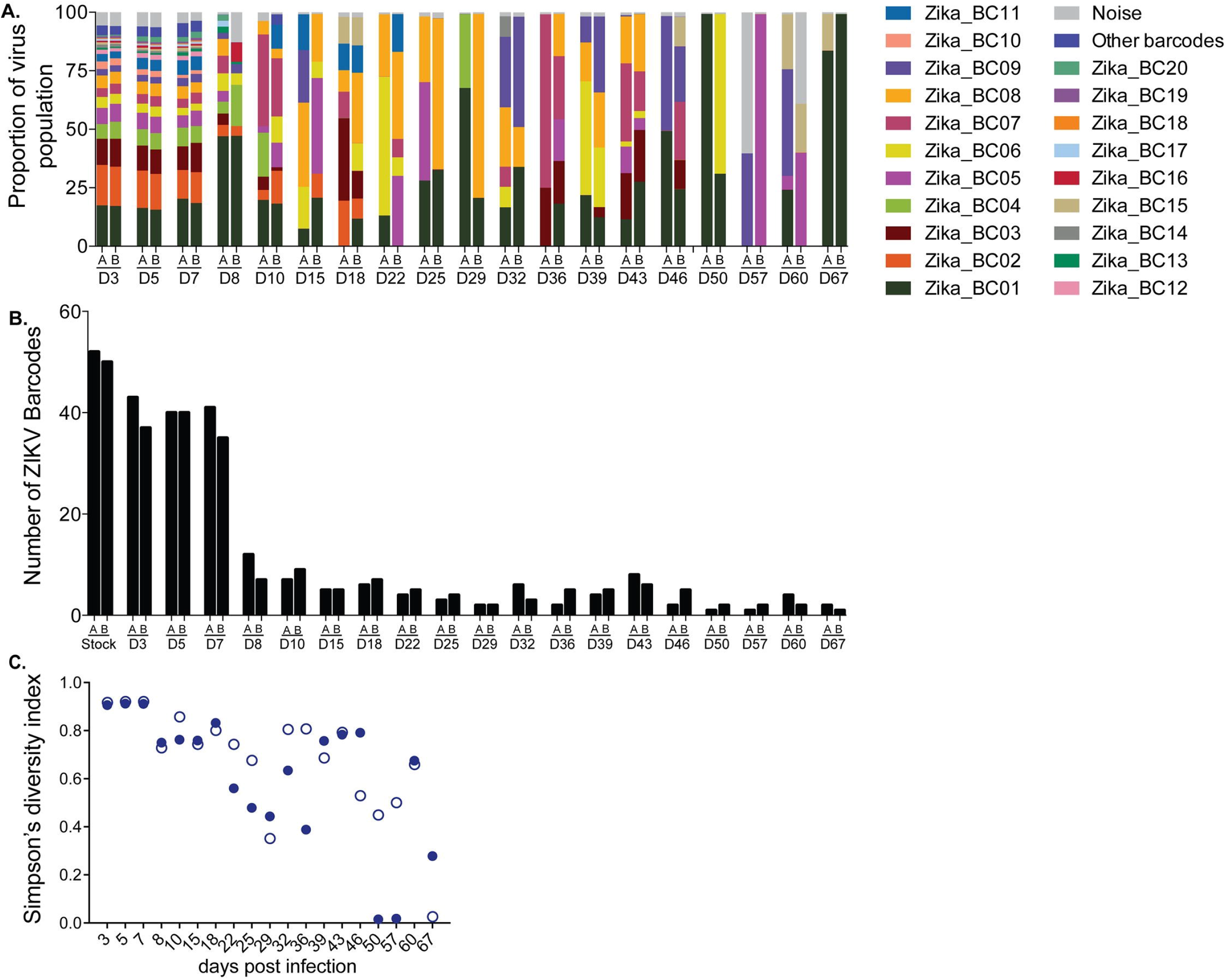
Sequencing of the molecular barcode isolated from pregnant animal 776301. Viral RNA was isolated from animal 776301 at the indicated time points. The theoretical number of cDNA molecules used in each PCR reaction is shown in Table 5. For each sample, a single preparation of cDNA was made and then split into two separate PCR reactions with primer set A (178 bp) and B (131bp). PCR products were tagged and sequenced. **A.**) The frequency of each barcode detected was quantified. ‘Other barcodes’ were the barcodes present in the list of the Top 57. ‘Noise’ represents sequences detected in the barcode region, but did not match the Top 57. **B.**) The number of barcodes detected at a frequency of greater than 0.047% in animal 776301 were counted for each sample and the data for each individual replicate are shown. **C.**) Average genetic complexity at the barcode positions measured by Simpson’s diversity index. Closed symbols represent Simpson’s diversity index in replicate A samples and open symbols represent Simpson’s diversity index in replicate B.

**Table 5.**
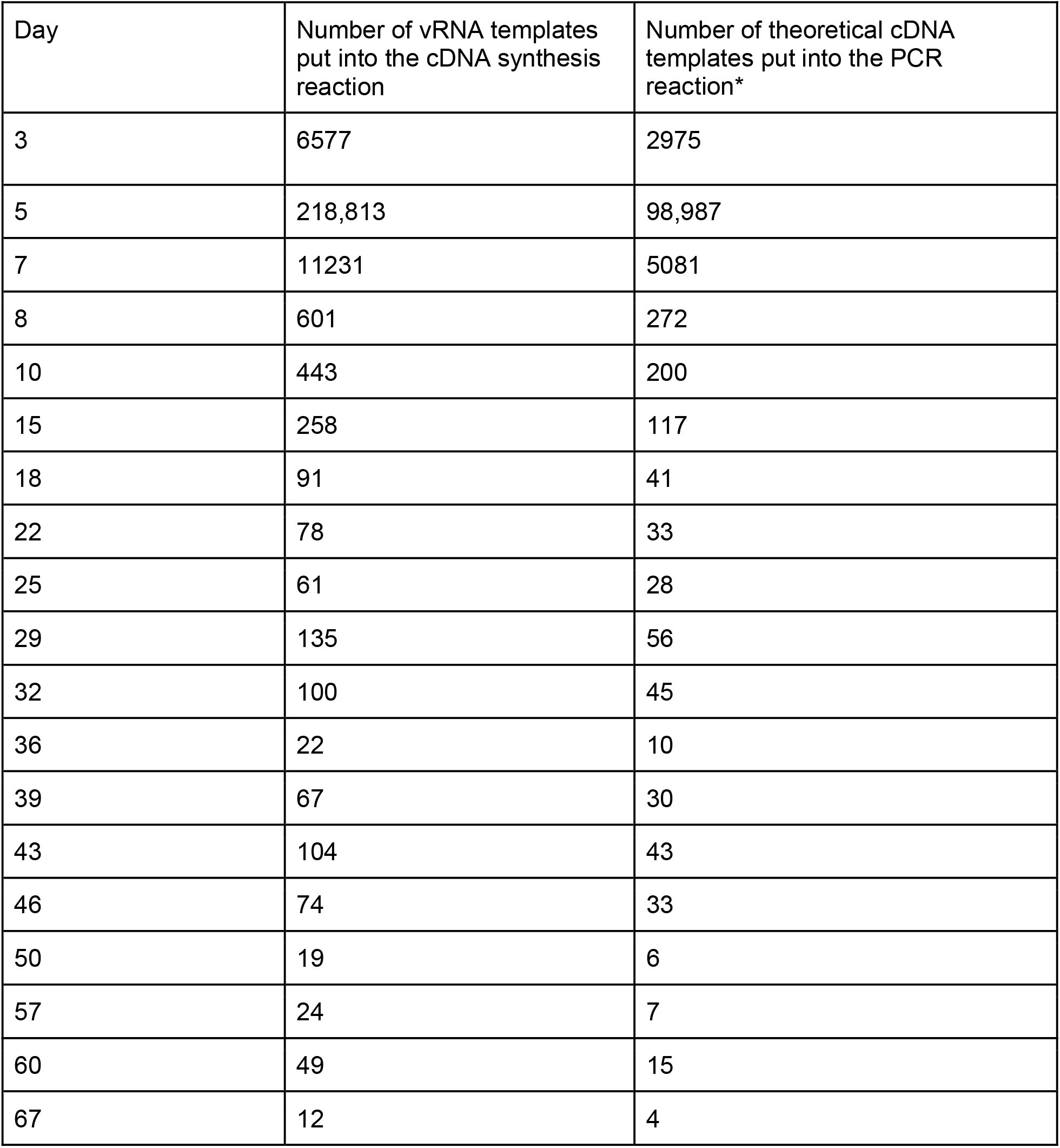
Number of viral templates sequenced from animal 776301 who was infected with ZIKV-BC-1.0

| Day | Number of vRNA templates put into the cDNA synthesis reaction | Number of theoretical cDNA templates put into the PCR reaction* |
| --- | --- | --- |
| 3 | 6577 | 2975 |
| 5 | 218,813 | 98,987 |
| 7 | 11231 | 5081 |
| 8 | 601 | 272 |
| 10 | 443 | 200 |
| 15 | 258 | 117 |
| 18 | 91 | 41 |
| 22 | 78 | 33 |
| 25 | 61 | 28 |
| 29 | 135 | 56 |
| 32 | 100 | 45 |
| 36 | 22 | 10 |
| 39 | 67 | 30 |
| 43 | 104 | 43 |
| 46 | 74 | 33 |
| 50 | 19 | 6 |
| 57 | 24 | 7 |
| 60 | 49 | 15 |
| 67 | 12 | 4 |

### Using ZIKV-BC-1.0 to evaluate transmission bottlenecks

To begin to understand potential transmission bottlenecks within the vector and the impact they might have on ZIKV population diversity, *Aedes aegypti* vector competence for ZIKV-BC-1.0 was evaluated at days 7, 13, and 25 days post feeding (PF) from mosquitoes that were exposed to the pregnant macaque at 4 dpi. A single *Ae. aegypti* was transmission-competent at day 25 PF (**Table 6**) as measured by plaque assay. Infection efficiency indicates the proportion of mosquitoes with virus-positive bodies among the tested ones. Dissemination efficiency indicates the proportion of mosquitoes with virus-positive legs, and transmission efficiency indicates the proportion of mosquitoes with infectious saliva among the tested ones. All other mosquitoes screened using this methodology were ZIKV-negative. These data are consistent with field epidemiological reports, which estimated mosquito infection rates during ZIKV outbreaks to be 0.061% [25] and also are consistent with infection rates during DENV and chikungunya outbreaks [26]. We also found low mosquito infection rates in a previous study exposing mosquitoes to ZIKV-infected rhesus macaques [22]. We deep sequenced virus (viral template numbers added to cDNA synthesis reactions are listed in **Table 7**) from all three anatomical compartments from this mosquito (body, leg, and saliva), and we only detected the presence of a single barcode: Zika_BC02. The viral loads in the body, leg, and saliva were 2.57 × 10^8^, 4.73 × 10^7^, and 4.29 × 10^4^ vRNA copies/ml, respectively. Zika_BC02 was present in the pregnant animal’s virus population at ~15% between days 3 and 5 after infection, representing the second most common barcode in the population (**Fig 6, Table 7, and Table S4**).

**Figure 6.**
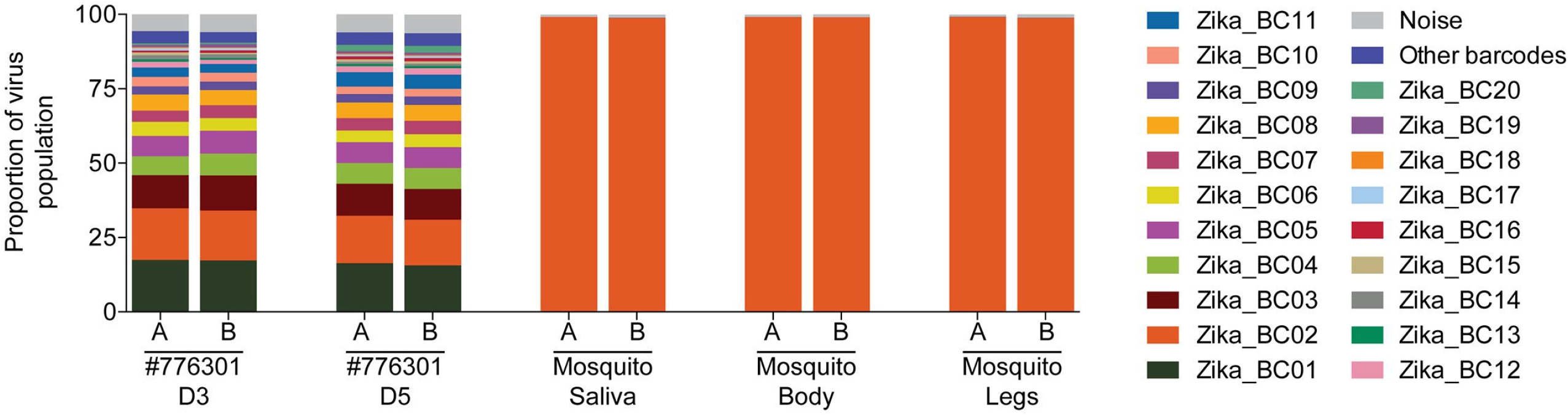
Sequencing of the molecular barcode isolated from mosquito #27 that fed on 776301. Viral RNA that was isolated from the saliva, body, and legs of mosquito #27 at day 25 post feeding was converted into cDNA and then split into two separate PCR reactions with primer set A (178bp) and B (131bp). The theoretical number of cDNA molecules used in each PCR reaction is shown in Table 7. PCR products were tagged and sequenced. The frequency of each barcode detected was quantified. Mosquito #27 fed on 776301 on day 4, so the frequencies of barcodes detected in the plasma of 776301 from figure 5 are shown. ‘Other barcodes’ were the barcodes present in the list of the Top 57. ‘Noise’ represents sequences detected in the barcode region, but did not match the Top 57.

**Table 6.** Vector competence of *Aedes aegypti* following peroral exposure to ZIKV-BC-1.0-infected pregnant macaque 4 days post inoculation.

| 7 days post feeding |  |  | 13 days post feeding |  |  | 25 days post feeding |  |  |
| --- | --- | --- | --- | --- | --- | --- | --- | --- |
| I | D | T | I | D | T | I | D | T |
| 0/30 (0%) | 0/30 (0%) | 0/30 (0%) | 0/30 (0%) | 0/30 (0%) | 0/30 (0%) | 1/30 (3%) | 1/30 (3%) | 1/30 (3%) |

I, % Infected D, % Disseminated T, % Transmitting

**Table 7.** Number of viral templates sequenced from one positive mosquito who fed on 776301

| Sample | Number of vRNA templates put into the cDNA synthesis reaction | Number of theoretical cDNA templates put into the PCR reaction* |
| --- | --- | --- |
| Mosquito saliva | 2830 | 1280 |
| Mosquito body | 10,000 | 4,524 |
| Mosquito legs | 10,000 | 4,524 |

The number of cDNA templates was calculated based on the number of vRNA templates that were put into the cDNA synthesis reaction, and then the amount of cDNA that was used for the PCR reaction.

## Discussion

Mosquito-borne viruses like ZIKV typically exist in hosts as diverse mutant swarms. Defining the way in which stochastic forces within hosts shape these swarms is critical to understanding the evolutionary and adaptive potential of these pathogens and may reveal key insight into transmission, pathogenesis, immune evasion, and reservoir establishment. To date, no attempts have been made to enumerate and characterize individual viral lineages during ZIKV infection. Here we characterized the dynamics of ZIKV infection in rhesus macaques and mosquitoes. Specifically, using a synthetic swarm of molecularly barcoded ZIKV, we tracked the composition of the virus population in mosquitoes and over time in both pregnant and nonpregnant animals.

Our results demonstrated that viral diversity fluctuated in both a spatial and temporal manner as host barriers or selective pressures were encountered and this likely contributed to narrowing of the barcode composition in both macaques and mosquitoes. For example, the proportions of individual barcoded virus templates remained stable during acute infection, but in the pregnant animal infected with ZIKV-BC-1.0 the complexity of the virus population declined precipitously 8 days following infection of the dam. This was coincident with the timing of typical resolution of ZIKV in non-pregnant macaques (**Figs 2 and 3**), and after this point the complexity of the virus population remained low for the subsequent duration of viremia (**Fig 5C**). We speculate that the narrowing of the barcode composition in the pregnant animal was the result of establishment of an anatomic reservoir of ZIKV that is not accessible to maternal neutralizing antibodies, which is shed into maternal plasma at low, but detectable, levels. It also is possible that declining viral barcode diversity was an artifact of a declining viral population size and the consequent effects on sampling, without reservoir establishment. Unfortunately, the absence of ZIKV RNA in the fetus at term prevented us from comparing the barcode composition in the fetus to the barcodes in maternal plasma, so this experiment could not resolve questions related to the potential that the feto-placental unit acts as a tissue reservoir of ZIKV. Connecting ZIKV clonotypes in neonatal tissues with clonotypes found in the mother will be important for understanding vertical transmission. Nevertheless, we demonstrated that synthetic swarm viruses can be used to probe the composition of viral populations over time *in vivo* in both macaques and mosquitoes; such synthetic swarms will be useful tools for future studies aimed at understanding vertical transmission, persistent reservoirs, bottlenecks, and overall evolutionary dynamics. While the ZIKV-BC-1.0 reported here has limited complexity, we have recently developed a new synthetic swarm, ZIKV-BC-2.0, which uses an optimized transfection strategy and has orders of magnitude more putative authentic barcodes. This new virus will be used in future studies in conjunction with deep sequencing techniques that enumerate individual templates with unique molecular identifiers [27]. We therefore expect that future studies of pregnant animals infected with barcoded ZIKV will help distinguish between these possibilities.

Although we developed this system to better understand the dynamics of ZIKV infection in the vertebrate host, this approach can be applied to address other questions about ZIKV transmission. For example, ZIKV-BC-1.0 can be used to quantify the bottleneck forces during mosquito infection and transmission. As a result, we also attempted to characterize barcodes present in mosquitoes that fed on the ZIKV-BC-1.0-infected pregnant animal. Consistent with our previous experiments [22], only a single *Ae. aegypti* became infected with ZIKV-BC-1.0 after feeding on ZIKV-BC-1.0-viremic macaques. This was likely the result of the low amount of infectious virus in macaque blood [28]. We only detected a single barcode during infection of mosquitoes. This is not entirely surprising because mosquitoes ingest small amounts of blood from infected hosts, which limits the size of the viral population founding infection in the vector. For example it has been previously estimated that as few as 5-42 founder viruses initiate DENV infection of the mosquito midgut [29]. Also, during replication in mosquitoes, flaviviruses undergo population bottlenecks as they traverse physical barriers like the midgut and salivary glands [29,30]. We therefore expected barcode diversity to be low in infected mosquitoes and these data are perhaps indicative of a stringent midgut bottleneck in this individual that limited the variant pool in other anatomic compartments, but this requires further experimental confirmation. Consistent with what we show here, previous work has demonstrated considerable haplotype turnover for West Nile virus in *Culex pipiens* but not in *Ae. aegypti*, i.e., haplotypes remained relatively stable as the virus trafficked from the midgut to the saliva [30]. Likewise, **Weger-Lucarelli et al., concomitant submission** most often detected only a single barcode in different *Ae. aegypti* populations that were exposed to ZIKV-BC-1.0 using an artificial membrane feeding system. In sum, our approach showed that synthetic swarm viruses can be used to probe the composition of viral populations over time *in vivo* to understand vertical transmission, persistent reservoirs, bottlenecks, and evolutionary dynamics.

## Materials and Methods

### Study Design

This study was a proof of concept study designed to examine whether molecularly barcoded ZIKV could be used to elucidate the source of prolonged maternal viremia during pregnancy (**Fig 3**). Datasets used in this manuscript are publicly available at zika.labkey.com.

### Ethical approval

This study was approved by the University of Wisconsin-Madison Institutional Animal Care and Use Committee (Animal Care and Use Protocol Number G005401).

### Nonhuman primates

Five male and five female Indian-origin rhesus macaques utilized in this study were cared for by the staff at the Wisconsin National Primate Research Center (WNPRC) in accordance with the regulations, guidelines, and recommendations outlined in the Animal Welfare Act, the Guide for the Care and Use of Laboratory Animals, and the Weatherall report. In addition, all macaques utilized in the study were free of Macacine herpesvirus 1, Simian Retrovirus Type D, Simian T-lymphotropic virus Type 1, and Simian Immunodeficiency Virus. For all procedures, animals were anesthetized with an intramuscular dose of ketamine (10mL/kg). Blood samples were obtained using a vacutainer or needle and syringe from the femoral or saphenous vein. The pregnant animal (776301) had a previous history of experimental DENV-3 exposure, approximately one year prior to ZIKV infection.

### Cells and viruses

African Green Monkey kidney cells (Vero; ATCC #CCL-81) were maintained in Dulbecco’s modified Eagle medium (DMEM) supplemented with 10% fetal bovine serum (FBS; Hyclone, Logan, UT), 2 mM L-glutamine, 1.5 g/L sodium bicarbonate, 100 U/ml penicillin, 100 μg/ml of streptomycin, and incubated at 37°C in 5% CO_2_. *Aedes albopictus*mosquito cells were (C6/36; ATCC #CRL-1660) were maintained in DMEM supplemented with 10% fetal bovine serum (FBS; Hyclone, Logan, UT), 2 mM L-glutamine, 1.5 g/L sodium bicarbonate, 100 U/ml penicillin, 100 μg/ml of streptomycin, and incubated at 28°C in 5% CO_2_. ZIKV strain PRVABC59 (ZIKV-PR; GenBank:KU501215), originally isolated from a traveler to Puerto Rico with three rounds of amplification on Vero cells, was obtained from Brandy Russell (CDC, Ft. Collins, CO). Virus stocks were prepared by inoculation onto a confluent monolayer of C6/36 mosquito cells with two rounds of amplification. A single harvest with a titer of 1.58 x 10^7^ plaque forming units (PFU) per ml (equivalent to 2.01 x 10^10^ vRNA copies per ml) of Zika virus/H.sapiens-tc/PUR/2015/PRVABC59-v3c2 were used for challenges utilizing wild type virus. This virus also served as the backbone upon which the genetically-barcoded virus was generated.

### Construction of molecularly-barcoded ZIKV

Genetically-barcoded ZIKV was constructed using the ZIKV reverse genetic platform developed by Weger-Lucarelli et al. [21]. The region for the barcode insertion was selected by searching for consecutive codons in which inserting a degenerate nucleotide in the third position would result in a synonymous change. The genetically-barcoded ZIKV clone then was constructed using a novel method called bacteria-free cloning (BFC). First, the genome was amplified as two overlapping pieces from the two-part plasmid system of the reverse genetic platform (see [21]). The CMV promoter was amplified from pcDNA3.1 (Invitrogen). The barcode region was then introduced in the form of an overlapping PCR-amplified oligo (IDT, Iowa, USA). All PCR amplifications were performed with Q5 DNA polymerase (New England Biolabs, Ipswich, MA, USA). Amplified pieces were then gel purified (Macherey-Nagel). The purified overlapping pieces were then assembled using the HiFi DNA assembly master mix (New England Biolabs) and incubated at 50°C for four hours. The Gibson assembly reaction then was treated with Exonuclease I (specific for ssDNA), lambda exonuclease (removes non-circular dsDNA) and DpnI (removes any original bacteria derived plasmid DNA) at 37°C for 30 minutes followed by heat inactivation for 20 minutes at 80°C. Two microliters of this reaction then was used for rolling circle amplification (RCA) using the REPLI-g Mini kit (Qiagen). RCA was performed following the manufacturer’s specifications except that 2M trehalose was used in place of water in the reaction mixture because it has been previously shown that this modification reduces secondary amplification products [31]. Reactions were incubated at 30°C for four hours and then inactivated at 65°C for three minutes. Sequence was confirmed by sanger sequencing.

### Molecularly-barcoded ZIKV stocks

Virus was prepared in Vero cells transfected with the purified RCA reaction. Briefly, RCA reactions were digested with NruI at 37°C for one hour to linearize the product and remove the branched structure. Generation of an authentic 3’UTR was assured due to the presence of the hepatitis-delta ribozyme immediately following the viral genome [21]. The digested RCA reaction then was purified using a PCR purification kit (Macherey-Nagel) and eluted with molecular-grade water. Purified and digested RCAs were transfected into 80-90% confluent Vero cells using the Xfect transfection reagent (Clontech) following manufacturer’s specifications. Infectious virus was harvested when 50-75% cytopathic effects were observed (6 days post transfection). Viral supernatant then was clarified by centrifugation and supplemented to a final concentration of 20% fetal bovine serum and 10 mM HEPES prior to freezing and storage as single use aliquots. Titer was measured by plaque assay on Vero cells as described in a subsequent section.

### Subcutaneous inoculations

The ZIKV-PR stock, ZIKV-IC, and ZIKV-BC-1.0 were thawed, diluted in PBS to 1 x 10^4^ PFU/mL, and loaded into a 3mL syringe maintained on ice until inoculation. Each of nine nonpregnant Indian-origin rhesus macaques was anesthetized and inoculated subcutaneously over the cranial dorsum with 1mL ZIKV-PR stock (n=3), ZIKV-IC stock (n=3), or ZIKV-BC-1.0 stock (n=3) containing 1 x 10^4^ PFU. Likewise, the pregnant animal was anesthetized and inoculated via the same route with 1 mL barcoded virus stock containing 1 x 10^4^ PFU. All animals were closely monitored by veterinary and animal care staff for adverse reactions and signs of disease. Nonpregnant animals were examined, and blood and urine were collected from each animal daily from 1 through 10 days, and 14 days post inoculation (dpi). Sampling continued for the pregnant animal until the resolution of viremia.

### Mosquito strain, colony maintenance, and vector competence

The *Aedes aegypti* black-eyed Liverpool (LVP) strain used in this study was obtained from Lyric Bartholomay (University of Wisconsin-Madison, Madison, WI) and maintained at the University of Wisconsin-Madison as previously described [32]. *Ae. aegypti* LVP are susceptible to ZIKV [33]. Infection, dissemination, and transmission rates were determined for individual mosquitoes and sample sizes were chosen using long established procedures [33–35]. Mosquitoes that fed to repletion on macaques were randomized and separated into cartons in groups of 40-50 and maintained as described in a previous section. All samples were screened by plaque assay on Vero cells. Dissemination was indicated by virus-positive legs. Transmission was defined as release of infectious virus with salivary secretions, i.e., the potential to infect another host, and was indicated by virus-positive salivary secretions.

### Plaque assay

All ZIKV screens from mosquito tissue and titrations for virus quantification from virus stocks were completed by plaque assay on Vero cell cultures. Duplicate wells were infected with 0.1 □ ml aliquots from serial 10-fold dilutions in growth media and virus was adsorbed for one hour. Following incubation, the inoculum was removed, and monolayers were overlaid with 3□ml containing a 1:1 mixture of 1.2% oxoid agar and 2X DMEM (Gibco, Carlsbad, CA) with 10% (vol/vol) FBS and 2% (vol/vol) penicillin/streptomycin. Cells were incubated at 37□°C in 5% CO2 for four days for plaque development. Cell monolayers then were stained with 3□ml of overlay containing a 1:1 mixture of 1.2% oxoid agar and 2X DMEM with 2% (vol/vol) FBS, 2% (vol/vol) penicillin/streptomycin, and 0.33% neutral red (Gibco). Cells were incubated overnight at 37□°C and plaques were counted.

### Plaque reduction neutralization test (PRNT)

Macaque serum samples were screened for ZIKV and DENV neutralizing antibody utilizing a plaque reduction neutralization test (PRNT) on Vero cells as described in [36] against ZIKV-PR and DENV-3. Neutralization curves were generated using GraphPad Prism software. The resulting data were analyzed by non-linear regression to estimate the dilution of serum required to inhibit 50% and 90% of infection.

### Fetal Rhesus Amniocentesis

Under real-time ultrasound guidance, a 22 gauge, 3.5 inch Quincke spinal needle was inserted into the amniotic sac. After 1.5-2 mL of fluid were removed and discarded due to potential maternal contamination, an additional 3-4 mL of amniotic fluid were removed for viral qRT-PCR analysis as described elsewhere [2,13]. These samples were obtained at the gestational ages 57, 71, 85, and 155 days. All fluids were free of any blood contamination.

### Viral RNA isolation

Plasma was isolated from EDTA-anticoagulated whole blood collected the same day by Ficoll density centrifugation at 1860 rcf for 30 minutes. Plasma was removed to a clean 15mL conical tube and centrifuged at 670 rcf for an additional 8 minutes to remove residual cells. Viral RNA was extracted from 300μL plasma using the Viral Total Nucleic Acid Kit (Promega, Madison, WI) on a Maxwell 16 MDx instrument (Promega, Madison, WI). Tissues were processed with RNAlater^®^ (Invitrogen, Carlsbad, CA) according to the manufacturer’s protocols. Viral RNA was isolated from the tissues using the Maxwell 16 LEV simplyRNA Tissue Kit (Promega, Madison, WI) on a Maxwell 16 MDx instrument. A range of 20-40 mg of each tissue was homogenized using homogenization buffer from the Maxwell 16 LEV simplyRNA Tissue Kit, the TissueLyser (Qiagen, Hilden, Germany) and two 5 mm stainless steel beads (Qiagen, Hilden, Germany) in a 2 ml snap-cap tube, shaking twice for 3 minutes at 20 Hz each side. The isolation was continued according to the Maxwell 16 LEV simplyRNA Tissue Kit protocol, and samples were eluted into 50 μl RNase free water. RNA was then quantified using quantitative RT-PCR. If a tissue was negative by this method, a duplicate tissue sample was extracted using the Trizol^TM^ Plus RNA Purification kit (Invitrogen, Carlsbad, CA). Because this purification kit allows for more than twice the weight of tissue starting material, there is an increased likelihood of detecting vRNA in tissues with low viral loads. RNA then was requantified using the same quantitative RT-PCR assay. Viral load data from plasma are expressed as vRNA copies/mL. Viral load data from tissues are expressed as vRNA/mg tissue.

### Cesarean section and tissue collection (Necropsy)

At ~155 days gestation, the fetus was removed via surgical uterotomy and maternal tissues were biopsied during laparotomy. These were survival surgeries for the dams. The entire conceptus (fetus, placenta, fetal membranes, umbilical cord, and amniotic fluid) was collected and submitted for necropsy. The fetus was euthanized with an overdose of sodium pentobarbitol (50 mg/kg). Tissues were dissected using sterile instruments that were changed between each organ and tissue type to minimize possible cross contamination. Each organ/tissue was evaluated grossly *in situ*, removed with sterile instruments, placed in a sterile culture dish, and sectioned for histology, viral burden assay, or banked for future assays. Sampling priority for small or limited fetal tissue volumes (e.g., thyroid gland, eyes) was vRNA followed by histopathology, so not all tissues were available for both analyses. Sampling of all major organ systems and associated biological samples included the CNS (brain, spinal cord, eyes), digestive, urogenital, endocrine, musculoskeletal, cardiovascular, hematopoietic, and respiratory systems as well as amniotic fluid, gastric fluid, bile, and urine. A comprehensive listing of all specific tissues collected and analyzed is presented in **Table 3**.

Biopsies of the placental bed (uterine placental attachment site containing deep decidua basalis and myometrium), maternal liver, spleen, and a mesenteric lymph node were collected aseptically during surgery into sterile petri dishes, weighed, and further processed for viral burden and when sufficient sample size was obtained, histology. Maternal decidua was dissected from the maternal surface of the placenta.

### Histology

Tissues (except neural tissues) were fixed in 4% paraformaldehyde for 24 hours and transferred into 70% ethanol until alcohol processed and embedded in paraffin. Neural tissues were fixed in 10% neutral buffered formalin for 14 days until routinely processed and embedded in paraffin. Paraffin sections (5μm) were stained with hematoxylin and eosin (H&E). Pathologists were blinded to vRNA findings when tissue sections were evaluated microscopically. Photomicrographs were obtained using a bright light microscope Olympus BX43 and Olympus BX46 (Olympus Inc., Center Valley, PA) with attached Olympus DP72 digital camera (Olympus Inc.) and Spot Flex 152 64 Mp camera (Spot Imaging), and captured using commercially available image-analysis software (cellSens DimensionR, Olympus Inc. and spot software 5.2).

### Quantitative reverse transcription PCR (qRT-PCR)

For ZIKV-PR, vRNA from plasma and tissues was quantified by qRT-PCR using primers with a slight modification to those described by Lanciotti et al. to accommodate African lineage ZIKV sequences [37]. The modified primer sequences are: forward 5’-CGYTGCCCAACACAAGG-3’, reverse 5’-CACYAAYGTTCTTTTGCABACAT-3’, and probe 5’-6fam-AGCCTACCTTGAYAAGCARTCAGACACYCAA-BHQ1-3’. The RT-PCR was performed using the SuperScript III Platinum One-Step Quantitative RT-PCR system (Invitrogen, Carlsbad, CA) on a LightCycler 480 instrument (Roche Diagnostics, Indianapolis, IN). The primers and probe were used at final concentrations of 600 nm and 100 nm respectively, along with 150 ng random primers (Promega, Madison, WI). Cycling conditions were as follows: 37°C for 15 min, 50°C for 30 min and 95°C for 2 min, followed by 50 cycles of 95°C for 15 sec and 60°C for 1 min. Viral RNA concentration was determined by interpolation onto an internal standard curve composed of seven 10-fold serial dilutions of a synthetic ZIKV RNA fragment based on a ZIKV strain derived from French Polynesia that shares >99% similarity at the nucleotide level to the Puerto Rican strain used in the infections described in this manuscript.

### Deep sequencing

Virus populations replicating in macaque plasma or mosquito tissues were sequenced in duplicate using a method adapted from Quick et. al. [38]. Viral RNA was isolated from mosquito tissues or plasma using the Maxwell 16 Total Viral Nucleic Acid Purification kit, according to manufacturer’s protocol. Viral RNA then was subjected to RT-PCR using the SuperScript IV Reverse Transcriptase enzyme (Invitrogen, Carlsbad, CA). Theoretical input viral template numbers are shown in Tables 2 to 5. For sequencing the entire ZIKV genome, the cDNA was split into two multi-plex PCR reactions using the PCR primers described in Quick et. al with the Q5^®^ High-Fidelity DNA Polymerase enzyme (New England Biolabs®, Inc., Ipswich, MA). For sequencing solely the barcode region, individual PCR reactions were performed that either used a primer pair generating a 131bp amplicon (131F: 5’-TGGTTGGCAATACGAGCGATGGTT-3’; 131R: 5’-CCCCCGCAAGTAGCAAGGCCTG-3’) or a 178bp amplicon (178F: 5’-CCTTGGAAGGCGACCTGATGGTTCT-3’; 178R (same as 131R): 5’-CCCCCGCAAGTAGCAAGGCCTG-3’). Purified PCR products were tagged with the Illumina TruSeq Nano HT kit or the and sequenced with a 2 x 300 kit on an Illumina MiSeq.

### Sequence analysis

Amplicon data were analyzed using a workflow we term “Zequencer 2017”(https://bitbucket.org/dhoconno/zequencer/src). Briefly, sequences were analyzed using a series of custom scripts generated in Python, as follows: to characterize the entire ZIKV genome, up to 1000 reads spanning each of the 35 amplicons were extracted from the data set and then mapped to the Zika reference for PRVABC59 (Genbank:KU501215). Variant nucleotides were called using SNPeff [39], using a 5% cutoff. Mapped reads and reference scaffolds were loaded into Geneious Pro (Biomatters, Ltd., Auckland, New Zealand) for intrasample variant calling and differences between each sample and the KU501215 reference were determined. Sequence alignments of the stock viruses can be found in the sequence read archive: ZIKV-IC (Acc #SRX3258286); ZIKV-BC-1.0 (Acc #SRX3258287).

To characterize the barcodes and their frequencies, the 24 nucleotide barcodes were first extracted from the alignment. Then, identical duplicate barcodes were counted using ‘Find duplicates’ in Geneious, and FASTA files were exported. Custom python scripts were then used to convert the lists of barcodes to TSV files, and then pivot tables were used in Excel to quantify the frequency of each barcode in an animal at a given time point.

### Diversity and similarity analysis

The diversities of the sequence populations were evaluated using the Simpson’s diversity index:

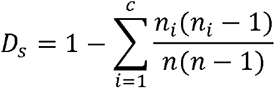

where *n_i_* is the number of copies of the *i*th unique sequence, *c* is the number of different unique sequences, and *n* is the total number of sequences in the sample.

The similarities between pairs of samples were assessed using the Morisita-Horn similarity index:

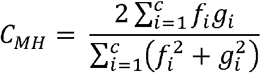

where *f_i_*=*n_1i_*/*N_1_* and *g_i_*=*n_2i_*/*N_2_*, *n_1i_* and *n_2i_* are the number of copies of the ith unique sequence in samples 1 and 2, and *N_1_* and *N_2_* are the total number of sequences in samples 1 and 2, respectively. The summations in the numerator and the denominator are over the *c* unique sequences in both samples.

The Simpson’s diversity and Morisita-Horn similarity indices account for both the number of unique sequences and their relative frequencies. These relative diversity and similarity indices range in value from 0 (minimal diversity/similarity) to 1 (maximal diversity/similarity). The Simpson’s diversity index considers a more diverse population as one with a more even distribution of sequence frequencies and the Morisita-Horn similarity index considers populations to be more similar if the higher frequency sequences in both samples are common to both samples and have similar relative frequencies. The diversity and similarity analyses were performed using Matlab (The Mathworks, Natick, MA).

### Data availability

Primary data that support the findings of this study are available at the Zika Open-Research Portal (https://zika.labkey.com). The authors declare that all other data supporting the findings of this study are available within the article and its supplementary information files, or from the corresponding author upon request.

## Acknowledgements

The authors acknowledge Jens Kuhn and Jiro Wada for preparing silhouettes of macaques used in figures. We thank the Veterinary, Animal Care, Scientific Protocol Implementation, and the Pathology staff at the Wisconsin National Primate Research Center (WNPRC) for their contribution to this study. We also thank Amy Ellis and Ryan Moriarty for sequencing efforts.

## Supporting Information

**Figure S1. Higher diversity is associated with higher number of input templates.** Sequence diversity vs. number of input templates (LHS) and total number of sequences per sample (RHS). The diversity measures include: number of unique sequences (upper) and Simpson’s diversity index (lower). The diversity for 3 replicate samples per input template number are shown.

**Figure S2. Higher similarity between replicate samples with higher number of input templates.** Similarity between pairs of replicate samples with the same input template number vs. number of input templates (LHS) and average total number of sequences (averaged between sample pairs) (RHS). The similarity measures include: number of common unique sequences (upper) and Morisita-Horn index (lower). The similarity between pairs of replicate samples per input template number are shown (i.e. RepA/RepB, RepA/RepC, and RepB/RepC).

**Figure S3. Higher similarity between sample pairs when both samples have a higher number of input templates.** Morisita-Horn similarity index between pairs of replicate samples with different input template numbers vs. number of input templates for the pair (i.e. sample 1/sample 2).

## References

1. Reynolds MR, Jones AM, Petersen EE, Lee EH, Rice ME, Bingham A et al. (2017) Vital Signs: Update on Zika Virus-Associated Birth Defects and Evaluation of All U.S. Infants with Congenital Zika Virus Exposure - U.S. Zika Pregnancy Registry, 2016. MMWR Morb Mortal Wkly Rep 66: 366–373.

2. Nguyen SM, Antony KM, Dudley DM, Kohn S, Simmons HA, Wolfe B et al. (2017) Highly efficient maternal-fetal Zika virus transmission in pregnant rhesus macaques. PLoS Pathog 13: e1006378.

3. Rosenberg K (2017) Zika Virus can Persist in Body Fluids for Prolonged Periods. Am J Nurs 117: 71.

4. Musso D, Roche C, Robin E, Nhan T, Teissier A, Cao-Lormeau VM (2015) Potential sexual transmission of Zika virus. Emerg Infect Dis 21: 359–361.

5. Aid M, Abbink P, Larocca RA, Boyd M, Nityanandam R, Nanayakkara O et al. (2017) Zika Virus Persistence in the Central Nervous System and Lymph Nodes of Rhesus Monkeys. Cell 169: 610–620.e14.

6. Hirsch AJ, Smith JL, Haese NN, Broeckel RM, Parkins CJ, Kreklywich C et al. (2017) Correction: Zika Virus infection of rhesus macaques leads to viral persistence in multiple tissues. PLoS Pathog 13: e1006317.

7. Prisant N, Bujan L, Benichou H, Hayot PH, Pavili L, Lurel S et al. (2016) Zika virus in the female genital tract. Lancet Infect Dis 16: 1000–1001.

8. Driggers RW, Ho CY, Korhonen EM, Kuivanen S, Jääskeläinen AJ, Smura T et al. (2016) Zika Virus Infection with Prolonged Maternal Viremia and Fetal Brain Abnormalities. N Engl J Med 374: 2142–2151.

9. Suy A, Sulleiro E, Rodó C, Vázquez é, Bocanegra C, Molina I et al. (2016) Prolonged Zika Virus Viremia during Pregnancy. N Engl J Med 375: 2611–2613.

10. Oliveira DB, Almeida FJ, Durigon EL, Mendes ÉA, Braconi CT, Marchetti I et al. (2016) Prolonged Shedding of Zika Virus Associated with Congenital Infection. N Engl J Med 375: 1202–1204.

11. Baud D, Van Mieghem T, Musso D, Truttmann AC, Panchaud A, Vouga M (2016) Clinical management of pregnant women exposed to Zika virus. Lancet Infect Dis 16: 523.

12. Aliota MT, Dudley DM, Newman CM, Mohr EL, Gellerup DD, Breitbach ME et al. (2016) Heterologous Protection against Asian Zika Virus Challenge in Rhesus Macaques. PLoS Negl Trop Dis 10: e0005168.

13. Dudley DM, Aliota MT, Mohr EL, Weiler AM, Lehrer-Brey G, Weisgrau KL et al. (2016) A rhesus macaque model of Asian-lineage Zika virus infection. Nat Commun 7: 12204.

14. Fennessey CM, Pinkevych M, Immonen TT, Reynaldi A, Venturi V, Nadella P et al. (2017) Genetically-barcoded SIV facilitates enumeration of rebound variants and estimation of reactivation rates in nonhuman primates following interruption of suppressive antiretroviral therapy. PLoS Pathog 13: e1006359.

15. Wu G, Webby RJ (2014) Barcoding influenza virus to decode transmission. Cell Host Microbe 16: 559–561.

16. Lauring AS, Andino R (2011) Exploring the fitness landscape of an RNA virus by using a universal barcode microarray. J Virol 85: 3780–3791.

17. Forrester NL, Guerbois M, Seymour RL, Spratt H, Weaver SC (2012) Vector-borne transmission imposes a severe bottleneck on an RNA virus population. PLoS Pathog 8: e1002897.

18. Pfeiffer JK, Kirkegaard K (2006) Bottleneck-mediated quasispecies restriction during spread of an RNA virus from inoculation site to brain. Proc Natl Acad Sci U S A 103: 5520–5525.

19. Varble A, Albrecht RA, Backes S, Crumiller M, Bouvier NM, Sachs D et al. (2014) Influenza A virus transmission bottlenecks are defined by infection route and recipient host. Cell Host Microbe 16: 691–700.

20. Ciota AT, Ehrbar DJ, Van Slyke GA, Payne AF, Willsey GG, Viscio RE et al. (2012) Quantification of intrahost bottlenecks of West Nile virus in Culex pipiens mosquitoes using an artificial mutant swarm. Infect Genet Evol 12: 557–564.

21. Weger-Lucarelli J, Duggal NK, Bullard-Feibelman K, Veselinovic M, Romo H, Nguyen C et al. (2017) Development and Characterization of Recombinant Virus Generated from a New World Zika Virus Infectious Clone. J Virol 91:

22. Dudley DM, Newman CM, Lalli J, Stewart LM, Koenig MR, Weiler AM et al. (2017) Infection via mosquito bite alters Zika virus tissue tropism and replication kinetics in rhesus macaques. Nature Communications

23. Newman CM, Dudley DM, Aliota MT, Weiler AM, Barry GL, Mohns MS et al. (2017) Oropharyngeal mucosal transmission of Zika virus in rhesus macaques. Nat Commun 8: 169.

24. Meaney-Delman D, Oduyebo T, Polen KN, White JL, Bingham AM, Slavinski SA et al. (2016) Prolonged Detection of Zika Virus RNA in Pregnant Women. Obstet Gynecol 128: 724–730.

25. Grubaugh ND, Ladner JT, Kraemer MUG, Dudas G, Tan AL, Gangavarapu K et al. (2017) Genomic epidemiology reveals multiple introductions of Zika virus into the United States. Nature 546: 401–405.

26. Dzul-Manzanilla F, Martínez NE, Cruz-Nolasco M, Gutierréz-Castro C, López-Damian L, Ibarra-López J et al. (2016) Evidence of vertical transmission and co-circulation of chikungunya and dengue viruses in field populations of Aedes aegypti (L.) from Guerrero, Mexico. Trans R Soc Trop Med Hyg 110: 141–144.

27. Zhou S, Jones C, Mieczkowski P, Swanstrom R (2015) Primer ID Validates Template Sampling Depth and Greatly Reduces the Error Rate of Next-Generation Sequencing of HIV-1 Genomic RNA Populations. J Virol 89: 8540–8555.

28. Ciota AT, Bialosuknia SM, Zink SD, Brecher M, Ehrbar DJ, Morrissette MN et al. (2017) Effects of Zika Virus Strain and Aedes Mosquito Species on Vector Competence. Emerg Infect Dis 23: 1110–1117.

29. Lequime S, Fontaine A, Ar Gouilh M, Moltini-Conclois I, Lambrechts L (2016) Genetic Drift, Purifying Selection and Vector Genotype Shape Dengue Virus Intra-host Genetic Diversity in Mosquitoes. PLoS Genet 12: e1006111.

30. Grubaugh ND, Weger-Lucarelli J, Murrieta RA, Fauver JR, Garcia-Luna SM, Prasad AN et al. (2016) Genetic Drift during Systemic Arbovirus Infection of Mosquito Vectors Leads to Decreased Relative Fitness during Host Switching. Cell Host Microbe 19: 481–492.

31. Pan X, Urban AE, Palejev D, Schulz V, Grubert F, Hu Y et al. (2008) A procedure for highly specific, sensitive, and unbiased whole-genome amplification. Proc Natl Acad Sci U S A 105: 15499–15504.

32. Christensen BM SDR (1984) Brugia pahangi: exsheathment and midgut penetration in Aedes aegypti. Transactions of the American Microscopical Society 103: 423–433.

33. Aliota MT, Peinado SA, Osorio JE, Bartholomay LC (2016) Culex pipiens and Aedes triseriatus Mosquito Susceptibility to Zika Virus. Emerg Infect Dis 22: 1857–1859.

34. Aliota MT, Peinado SA, Velez ID, Osorio JE (2016) The wMel strain of Wolbachia Reduces Transmission of Zika virus by Aedes aegypti. Sci Rep 6: 28792.

35. Aliota MT, Walker EC, Uribe Yepes A, Velez ID, Christensen BM, Osorio JE (2016) The wMel Strain of Wolbachia Reduces Transmission of Chikungunya Virus in Aedes aegypti. PLoS Negl Trop Dis 10: e0004677.

36. Lindsey HS, Calisher CH, Mathews JH (1976) Serum dilution neutralization test for California group virus identification and serology. J Clin Microbiol 4: 503–510.

37. Lanciotti RS, Kosoy OL, Laven JJ, Velez JO, Lambert AJ, Johnson AJ et al. (2008) Genetic and serologic properties of Zika virus associated with an epidemic, Yap State, Micronesia, 2007. Emerg Infect Dis 14: 1232–1239.

38. Quick J, Grubaugh ND, Pullan ST, Claro IM, Smith AD, Gangavarapu K et al. (2017) Multiplex PCR method for MinION and Illumina sequencing of Zika and other virus genomes directly from clinical samples. Nat Protoc 12: 1261–1276.

39. Cingolani P, Platts A, Wang LL, Coon M, Nguyen T, Wang L et al. (2012) A program for annotating and predicting the effects of single nucleotide polymorphisms, SnpEff: SNPs in the genome of Drosophila melanogaster strain w1118; iso-2; iso-3. Fly (Austin) 6: 80–92.

